# A transcription-centric model of SNP-Age interaction

**DOI:** 10.1101/2020.03.02.973388

**Authors:** Kun Wang, Mahashweta Basu, Justin Malin, Sridhar Hannenhalli

**Affiliations:** Cancer Data Science Lab, NCI, National Institutes of Health, Bethesda, MD; Center for Bioinformatics and Computational Biology, University of Maryland, College Park, MD; Laboratory of Genome Integrity, NCI, National Institutes of Health, Bethesda, MD

## Abstract

Complex age-associated phenotypes are caused, in part, by an interaction between an individual’s genotype and age. The mechanisms governing such interactions are however not entirely understood. Here, we provide a novel transcriptional mechanism-based framework – SNiPage, to investigate such interactions, whereby a transcription factor (TF) whose expression changes with age (age-associated TF), binds to a polymorphic regulatory element in an allele-dependent fashion, rendering the target gene’s expression dependent on both, the age and the genotype. Applying SNiPage to GTEx, we detected ~637 interacting TF-SNP-Gene triplets on average across 25 tissues, where the TF binds to a regulatory SNP in the gene’s promoter or putative enhancer and potentially regulates its expression in an age- and allele-dependent fashion. The detected SNPs are enriched for epigenomic marks indicative of regulatory activity, exhibit allele-specific chromatin accessibility, and spatial proximity to their putative gene targets. Furthermore, the TF-SNP interaction-dependent target genes have established links to aging and to age-associated diseases. In six hypertension-implicated tissues, detected interactions significantly inform hypertension state of an individual. Lastly, the age-interacting SNPs exhibit a greater proximity to the reported phenotype/diseases-associated SNPs than eSNPs identified in an interaction-independent fashion. Overall, we present a novel mechanism-based model, and a novel framework SNiPage, to identify functionally relevant SNP-age interactions in transcriptional control and illustrate their potential utility in understanding complex age-associated phenotypes.

## Introduction

Normal aging is a critical “environmental” risk factor for complex diseases such as hypertension, cardiovascular defects, macular degeneration, Parkinson’s disease, and cancers (Blasco et al., 2013; Niccoli and Partridge, 2012). For instance, in 2011-2014, the prevalence of hypertension among people older than 60 years was 65% compared to only 7.3% for people between 18 and 39 years old (Sug et al., 2015). Numerous crucial biological functions such as immune response, wound healing, DNA repair, metabolism, and mitochondria function. also significantly decline with aging (Blasco et al., 2013; Niccoli and Partridge, 2012; Wyss-coray, 2015), further underscoring aging as a major risk factor for complex diseases. Even though the underlying mechanisms linking aging with complex diseases are far from clear, profound transcriptomic changes associated with both aging and complex diseases have been identified (Glass et al., 2013; Tollervey et al., 2011; Wang et al., 2014a) and may provide mechanistic insights.

In addition to aging, genomic variations also significantly contribute to complex diseases (Adeyemo et al., 2009; Jiang et al., 2011; Visscher et al., 2017; Buniello et al., 2019), including cancer (Dror et al., 2016; Lubbe et al., 2012; Wang et al., 2014b). It is also widely accepted that genomic variations affect complex phenotypes, in substantial part, through transcriptomic variability. This is exemplified by numerous demonstrations that SNPs identified by eQTL studies (Acharya et al., 2017; Das et al., 2015; Gilad et al., 2008; Westra and Franke, 2014) are enriched in regulatory regions of the genome and for SNPs associated with complex traits, and these links have the potential to help identify key driver genes and mechanisms (Howard et al., 2019).

It is likely that aging and genome variations jointly contribute to complex diseases. That is, systemic molecular changes through aging may affect complex phenotypes in a genotype-dependent manner. Indeed, such genotype-environment (‘Age’ being an environmental factor in this scenario) interactions have been previously investigated. For instance, by incorporating a SNP-age term in a regression model, Yao et al identified 10 age-dependent eQTL SNPs (eSNPs) in Whole Blood (Yao et al, 2014). In a meta-analysis, Simino et al detected 9 SNPs which have age-dependent association with blood pressure (Simino et al., 2014). Work by Dongen et al showed that the methylome in whole blood could be affected by the interactions between SNPs and age (Dongen et al., 2016). However, these previous studies have not investigated specific molecular mechanisms underlying the SNP-age interactions in determining a phenotype, which limits biological insights they provide and, as will be made clear below, limits their statistical power as well.

Here, based on established transcriptional mechanisms, we present a model and a pipeline, that we term “SNiPage”, to identify SNP-Age interactions in determining target gene expression. Our model is based on the hypothesis that age-associated transcription factors (TF) exhibiting allele-specific binding at a regulatory SNP will lead to age-associated expression changes in the target gene in an allele-specific manner (Fig. 1). Thus, our model uses the expression level of an age-associated TF as a proxy for age and identifies TF-SNP-Gene triplets where the target gene’s expression is determined by an interaction between the TF’s expression level (equivalently, host age) and the SNP genotype. Our model thus facilitates exploration of SNP-age interactions across TFs and across tissues, substantially extending the scope and statistical power of previous explorations.

**Fig. 1.**
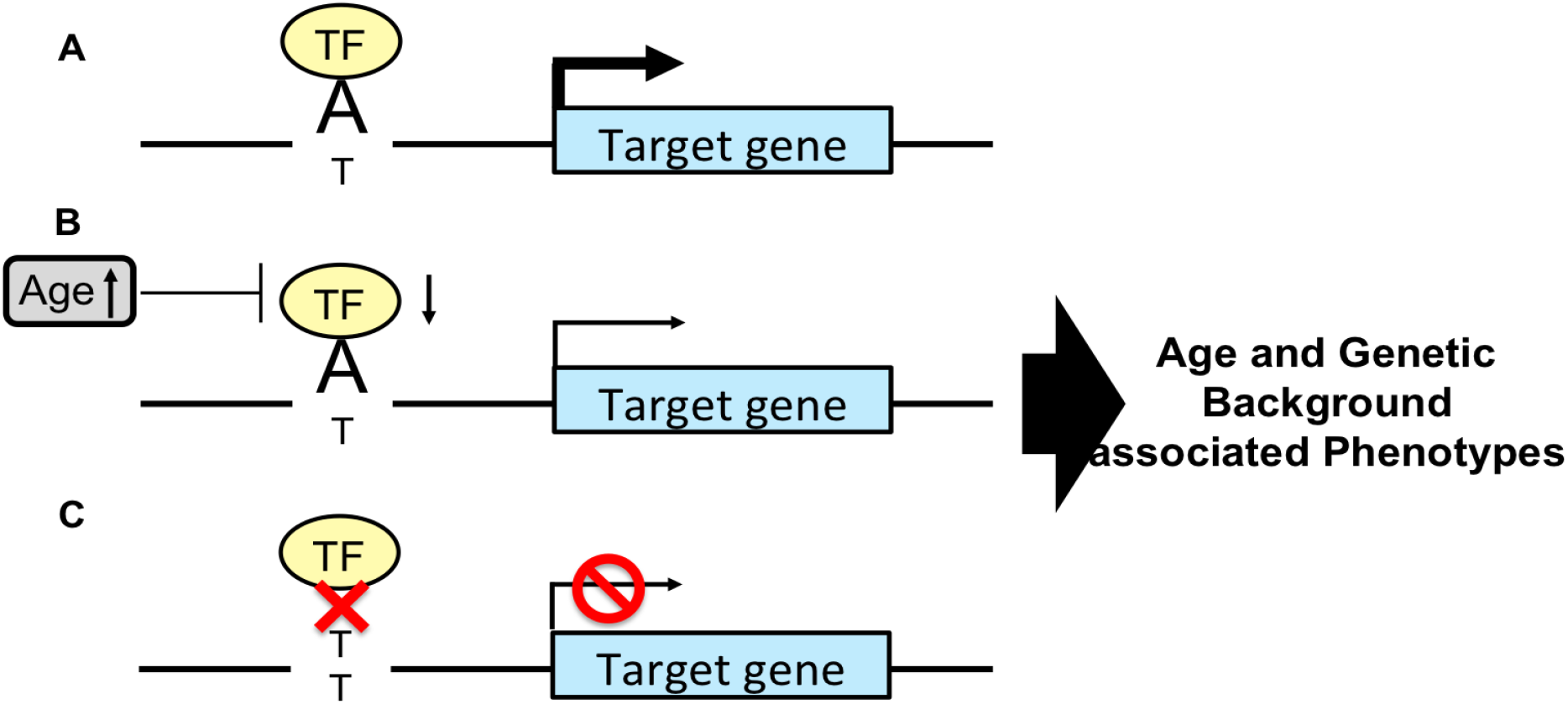
Hypothesized mechanism of SNP-Age interactions. (A) A TF binds to allele A to regulate the target gene. (B) With aging, the TF expression decreases and consequently so does the target gene’s expression for ‘A’ allele, but has not effect on the ‘T’ allele (C).

Application of SNiPage to 25 human tissues revealed 637 TF-SNP-Gene triplets on average in each tissue. Multiple biological evidence, including chromatin marks indicative of active regulation, allelic imbalance, and SNP-Gene spatial proximity, suggest that our detected SNP-Gene pairs are highly likely to be functional. Functional enrichment analyses link the detected target genes regulated by SNP-age interactions to the aging process and complex diseases. Interestingly, aggregated SNP-age interactions significantly inform an individual’s hypertension state in six known hypertension-related tissues. Likewise, for several other phenotypes, including cancers, the age-interacting SNPs exhibit a significantly higher linkage to the reported phenotype-associated SNPs than the SNPs detected using standard interaction-independent model. Finally, the detected SNPs exhibit relatively higher derived allele frequencies in human, suggestive of their potential role in adaptive evolution.

In summary, we have reported a novel framework, exploiting established transcriptional mechanisms, to identify SNP-age interactions driving gene expression, and the detected TF-SNP-Gene triplets may provide further insights into the mechanisms linking genotype and aging with complex age-related phenotypes.

## Results

### SNiPage overview

The SNiPage pipeline is illustrated in Fig.2 and described in Methods. We obtained the imputed SNP genotypes and transcriptome from ~570 individuals across 54 tissues from the GTEx consortium (GTEx Consortium, 2015) (sample details shown in supplementary Table 1). We retained those SNPs within open chromatin regions (using tissue-specific DNase hypersensitivity (DHS) (Roadmap Epigenomics Consortium et al., 2015)) which were within 1Mbp of gene promoters (Monlong et al., 2014; Rantalainen et al., 2015; Zhao et al., 2013). Our analysis is thus limited to those 25 tissues for which DHS profiles are available (supplementary Table 1). We detected age-associated TFs based on their gene expression (Methods), controlling for genetic background and hidden variables (Wang et al., 2014a; Wang et al., 2018; Yang et al., 2015). For age-associated TFs, based on published DNA-binding motifs, we identified their putative allele-specific binding at all retained SNPs (Methods). For each SNP, all genes within 1Mbp of the SNP are considered as its potential cis-regulatory targets. For each candidate TF-SNP-gene triplet, based on a linear model, we identified significant TF-SNP interactions associated with the target gene’s expression. Finally, as a further consistency filter, we selected only those TF-SNP-gene triplets that exhibited a TF-gene correlation only for the SNP allele promoting TF binding, and not for the other allele.

**Fig. 2.**
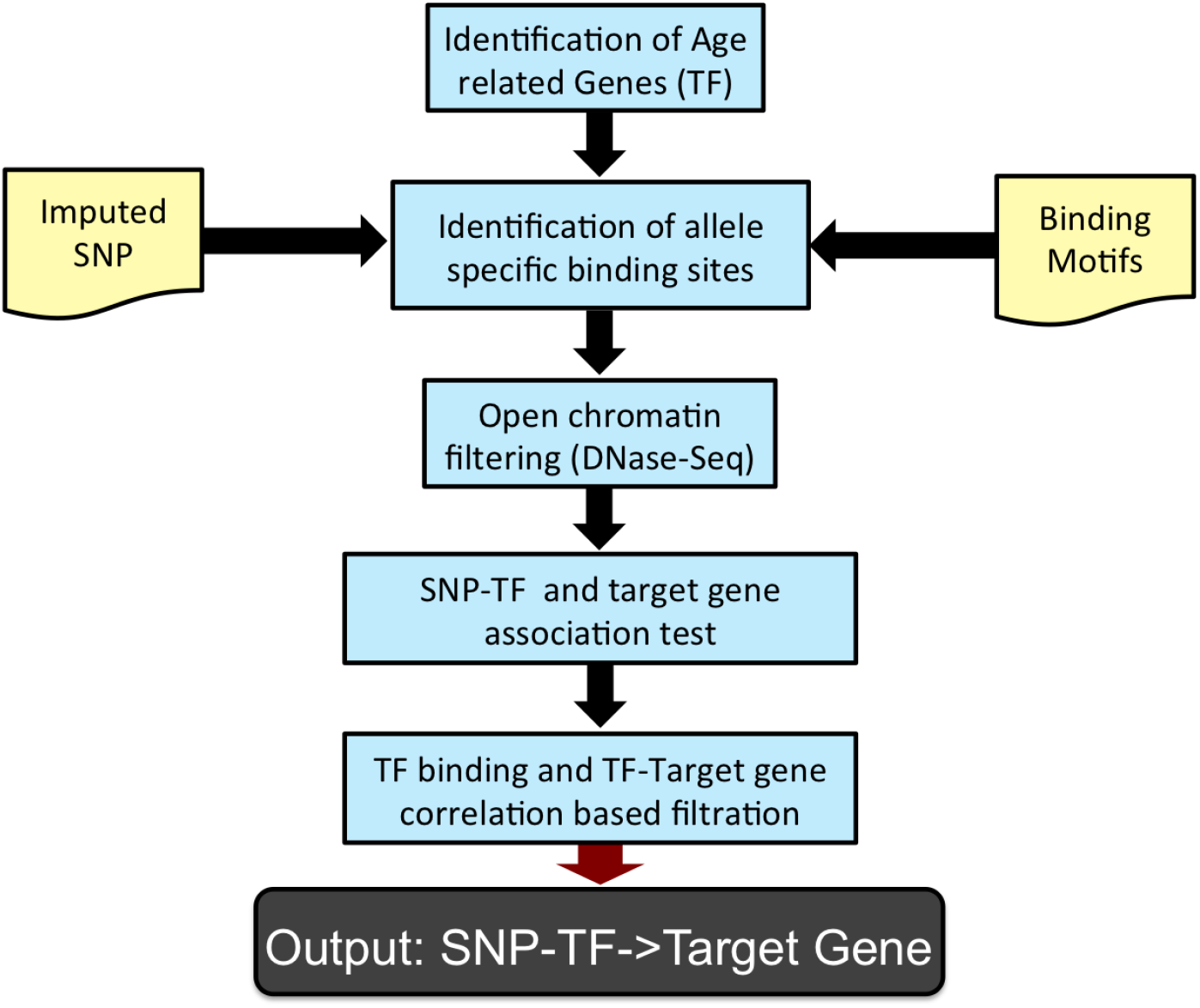
SNiPage pipeline. We identify age-associated TFs, and then identify genome-wide (imputed) SNPs at which those age-associated TFs are predicted to have allele-specific binding, based on their known DNA-binding motifs. The resulting SNPs are filtered to retain only those in open chromatin region, based on tissue-specific DNase-Seq data. Then we test the interaction significance using a linear model, which measures the association between the interaction and target gene expression. Finally, we further select the potential functional interactions based on consistency with allele-specific binding predictions and the TF-target gene expression correlations.

### SNiPage robustly detects numerous TF-SNP-Gene interactions across 25 tissues

We tested a total of 2,997,407 TF-SNP-gene triplets involving 125 TFs, 8,377 SNPs, and 4,661 genes on average across each of the 25 tissues and identified ~637 TF-SNP-Gene triplets (parameters at various steps of the pipeline are mentioned in Methods) on average in each tissue; details in Table 1. As a technical control, when we randomly shuffle gene expression across samples (permuting sample ids while preserving gene-gene covariance), far fewer triplets (~110 per tissue; Supplementary Table 2) are detected (p-value for paired Wilcoxon test for 25 tissues = 4.2e-07). We detected the largest number of interactions in whole blood, Brain-Cortex, and Colon-Transverse tissues. Interestingly, on average, only ~18% of the SNPs involved in a detected triplet were previously detected as eSNPs (GTEx Consortium, 2015). This potentially implies that numerous functional SNPs may escape detection based on standard interaction-independent eQTL studies due to interactions with environmental factors. On average only half (~53%) of the target genes involved in interactions are themselves significantly associated with age (enriched over the background control; Fisher test p-value = 5e-3), consistent with a regulatory role of age-associated TFs in driving age-associated expression of the target gene.

**Table 1.** The number of detected interaction triplets across 25 tissues.

| **Tissues** | **Number of triplets** | **Number of target genes** | **Number of TFs** | **Number of SNPs** |
| --- | --- | --- | --- | --- |
| Whole Blood | 2279 | 1220 | 212 | 1830 |
| Adipose-Subcutaneous | 165 | 149 | 81 | 160 |
| Muscle-Skeletal | 83 | 78 | 50 | 83 |
| Artery-Coronary | 50 | 48 | 13 | 49 |
| Heart-Atrial Appendage | 465 | 389 | 107 | 426 |
| Adipose-Visceral | 1403 | 1056 | 175 | 1244 |
| Ovary | 329 | 294 | 82 | 322 |
| Breast-Mammary | 536 | 458 | 143 | 504 |
| Brain-Cortex | 2080 | 1557 | 161 | 1827 |
| Adrenal Gland | 15 | 15 | 9 | 15 |
| Lung | 507 | 412 | 191 | 467 |
| Esophagus-Muscularis | 146 | 136 | 42 | 144 |
| Esophagus-Mucosa | 152 | 137 | 84 | 147 |
| Esophagus-Gastroesophageal Junction | 210 | 185 | 81 | 197 |
| Stomach | 169 | 158 | 47 | 168 |
| Colon-Sigmoid | 430 | 370 | 139 | 411 |
| Colon-Transverse | 2275 | 1475 | 128 | 1864 |
| Heart-Left Ventricle | 419 | 355 | 116 | 375 |
| Brain-Cerebellum | 20 | 19 | 6 | 20 |
| Artery-Aorta | 392 | 366 | 132 | 380 |
| Brain-Hippocampus | 932 | 777 | 60 | 862 |
| Brain-Frontal Cortex | 837 | 710 | 57 | 775 |
| Brain-Cerebellar Hemisphere | 1100 | 895 | 92 | 1014 |
| Brain-Caudate | 54 | 49 | 8 | 50 |
| Brain-Hypothalamus | 883 | 708 | 75 | 825 |

A specific example of a detected TF-SNP-Gene interaction is illustrated in Fig. 3A: TF *MAX* binds at SNP rs2295079 to regulate the expression of MTOR in Whole Blood. Expression of TF MAX significantly decreases with age (Fig. 3B). Notably, SNP rs2295079 was not captured by the GTEx eQTL study, even though its potential functional role is supported by DNase HS, H3K4me3, and H3K27ac peaks in K562, GM12878, and A549 tissues (Epigenetic regulation of RNA processing: Nature ENCODE: Nature Publishing Group). In addition, the *MAX* ChIP-Seq peak also appears in the same region in these cell lines, further supporting rs2295079’s functional role. Based on our putative allele-specific binding prediction for *MAX* at rs2295079, we expect it to differentially bind to the ‘G’ allele. Accordingly, we observed genotype-specific correlation between *MAX* and *MTOR*; as shown in Fig. 3C-E, *MTOR* expression is significantly negatively correlated (Spearman correlation = −0.24, p-value = 5.8e-3 for only one G allele; Spearman correlation = −0.3, p-value = 4e-4 for two G alleles) with *MAX* expression when the individuals have at least one major allele, while for ‘C’ allele homozygotes, there is no correlation between *MAX* and *MTOR* (Spearman Correlation = −0.03, p-value = 0.79).

**Fig. 3.**
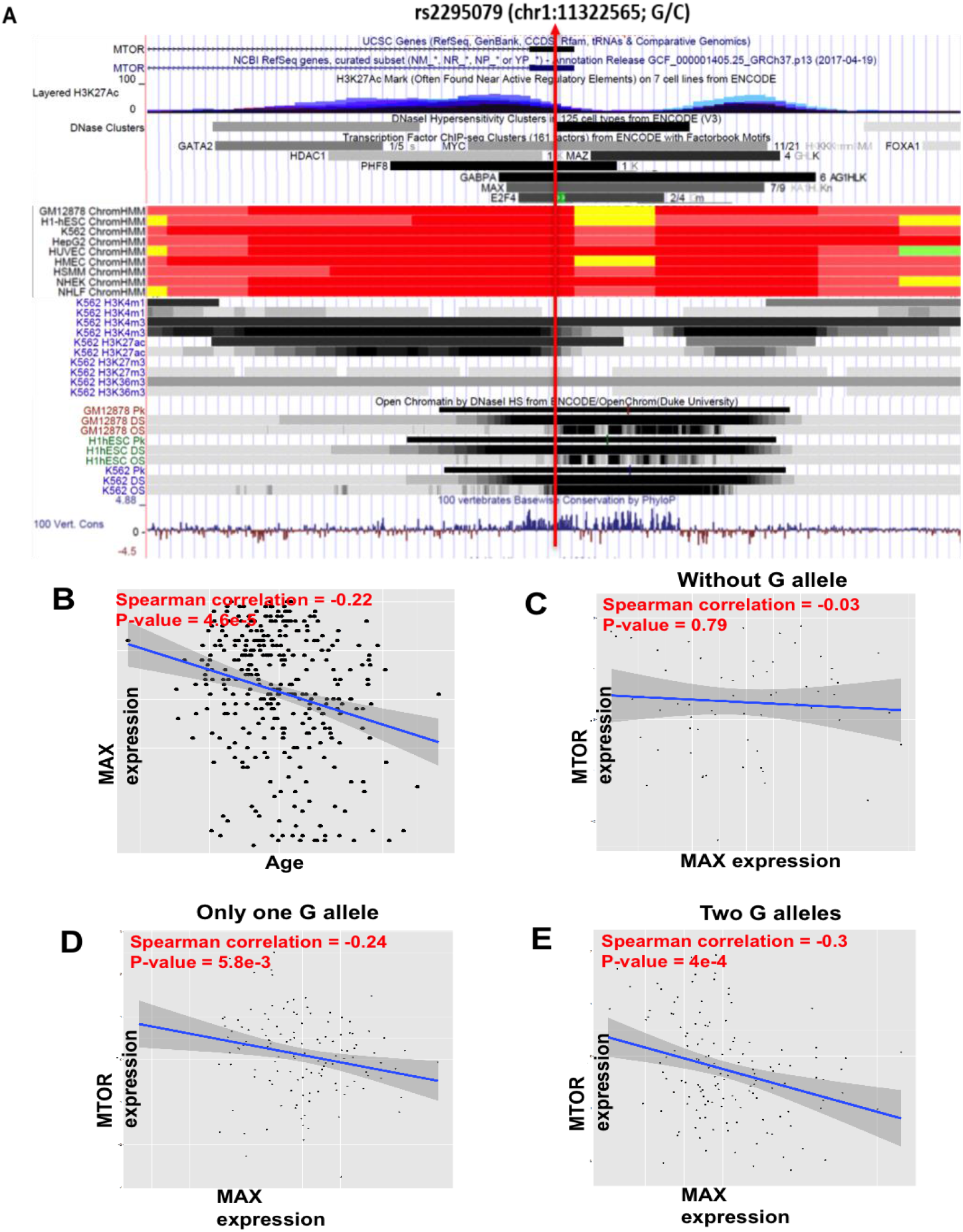
An example interacting TF-SNP-Gene triplet. (A) TF MAX binds to MTOR promoter region supported by DNase, H3K4me3, H3K27ac and MAX CHIP-Seq independently across several cell lines from ENCODE (K562, GM12878, and A549), and the binding is predicted to be specific to ‘G’ allele. (B) MAX gene is age-associated. (C)-(E) The correlations between MAX and MTOR across the three genotypes.

To assess the statistical robustness of the detected interactions, we estimated their replication rate in random down-sampled (90%, 80%, 70%) datasets in four tissues (adipose, whole blood, muscle and lung) with relatively large sample size. In addition, we also estimated the replication rate in the background dataset (randomly shuffled target gene expression profile) as control. As supplementary Fig. 1 shows, all under-sampled datasets across four tested tissues exhibit significantly higher replication rate than the background dataset (about 68%, 56%, 49% interaction triplets replicated respectively for 90%, 80%, 70% sampling datasets on average across 4 tissues in contrast to 0% replication rate for the background datasets). The decreasing trend of replication rates for 90%, 80%, 70% samples is consistent with the fact that sample size determine the statistical power. Overall, our analyses suggest that our detected interactions are not likely to be false positives (nominal FDR based on randomization ~ 17%) and are statistically robust.

### Identified interacting SNPs are likely to be functional

In view of our hypothesis that age-associated TFs bind to a SNP locus and regulate target gene expression, the detected SNP loci are expected to be in cis-regulatory elements such as enhancers and promoters. We therefore checked whether the detected SNP loci are enriched for open chromatin signals (DHS) and other histone marks (H3K27ac, H3K4me1 and H3K4me3) characterizing regulatory regions. Even though DHS peaks were used as an inclusion criterion for the SNPs, we tested whether the 100 bps flanks of the detected interacting SNPs exhibit higher DHS intensity compared to other tested SNPs which, notably, also qualified under the initial DHS filter. Each of the 4 epigenomic marks was analyzed for the tissues in which the relevant data was available and there were at least 100 foreground SNP loci. As shown in Fig. 4A, the foreground (SNP loci involved in the detected interactions) DHS intensities are significantly higher than the background (SNP loci that passed the initial DHS filter and were tested for interactions) in all 5 tissues analyzed. This was also broadly true for H3K27ac and H3K4me3 (Fig. 4C-D). However, we observed a significant difference between the foreground and the background SNPs for H3K4me1 in five of the fifteen tissues analyzed in Fig. 4B. Note that while H3K27ac and H3K4me1 are marker for active enhancers (Creyghton et al., 2010; Gibney and Nolan, 2010; Luco et al., 2010), H3K4me1 could also be related to poised enhancers (Creyghton et al., 2010). H3K4me3 is associated with gene promoters. Overall, these results strongly suggest that the SNPs involved in the detected interactions are likely to play a regulatory role.

**Fig. 4.**
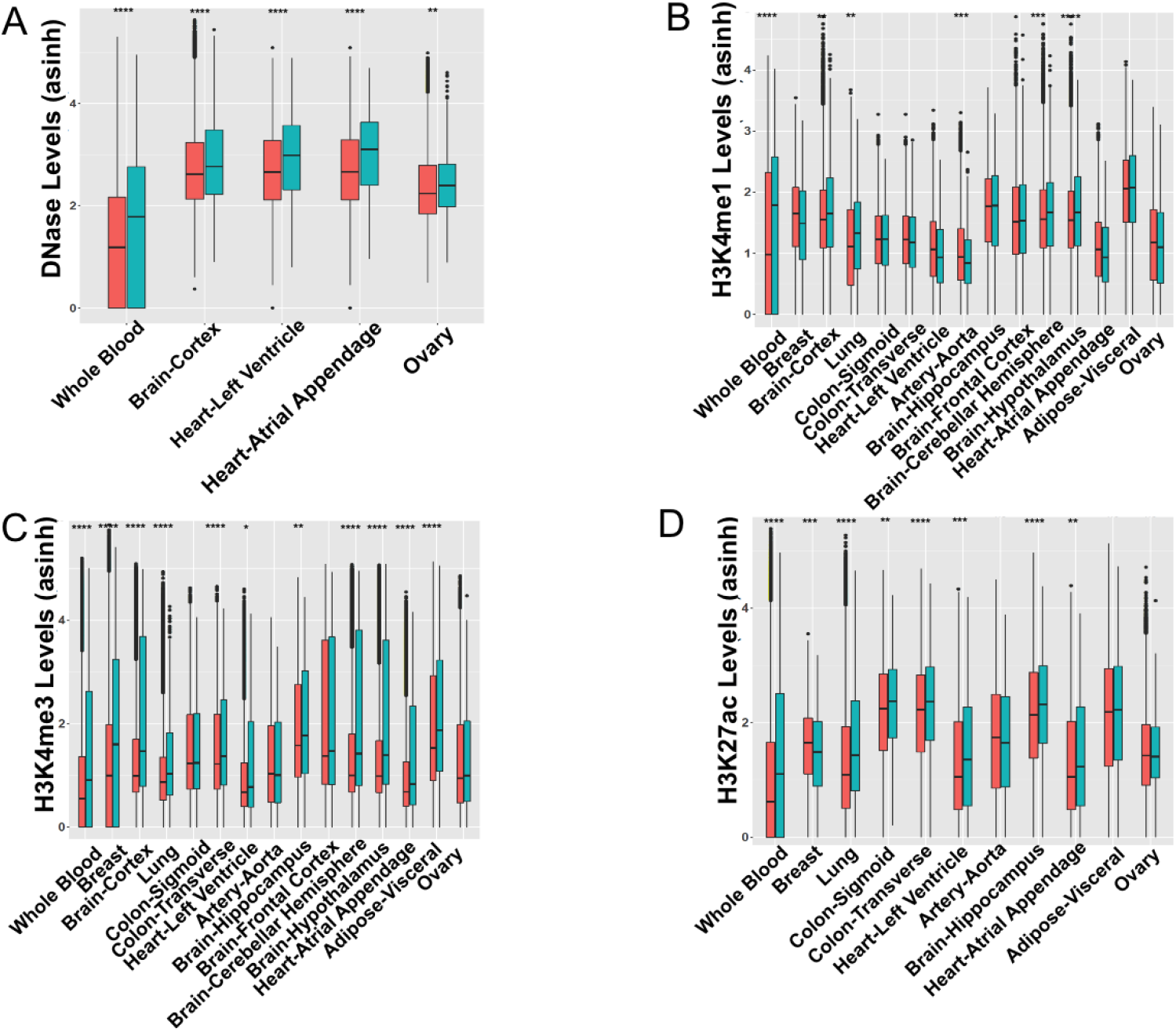
Detected SNPs are enriched for epigenomic markers of regulatory regions. (A)-(D) Epigenomic signal comparison between the foreground and control SNPs for open chromatin (DNase), H3K4me1, H3K4me3, and H3k27ac respectively. The Foreground is 100 bps windows around the detected SNPs (blue) and the control is 100bps windows around the other tested SNPs (red).

Next, we checked whether our detected SNPs are under evolutionary selection by comparing their conservation (PhastCons scores derived from 100 species multiple alignment) with those for three backgrounds: randomly selected SNPs, randomly selected SNPs after DNase filtration, and eSNPs. As shown in supplementary Fig. 2, our detected SNPs are significantly more conserved than random SNPs (in 21 tissues out of 25), but in fewer tissues relative to the latter two controls (in 14 and 6 tissues out of 25; supplementary Fig. 3-4). However, interestingly, our detected SNPs exhibit significantly higher derived allele frequency (DAF; see methods) (Auton et al., 2015) compared to eSNPs in 21 of the 25 tissues (Fig. 5A). These results suggest that SNPs that interact with age are likely to be in functional regions of the genome and are likely to be under positive or relaxed purifying selection during human evolution.

**Fig. 5.**
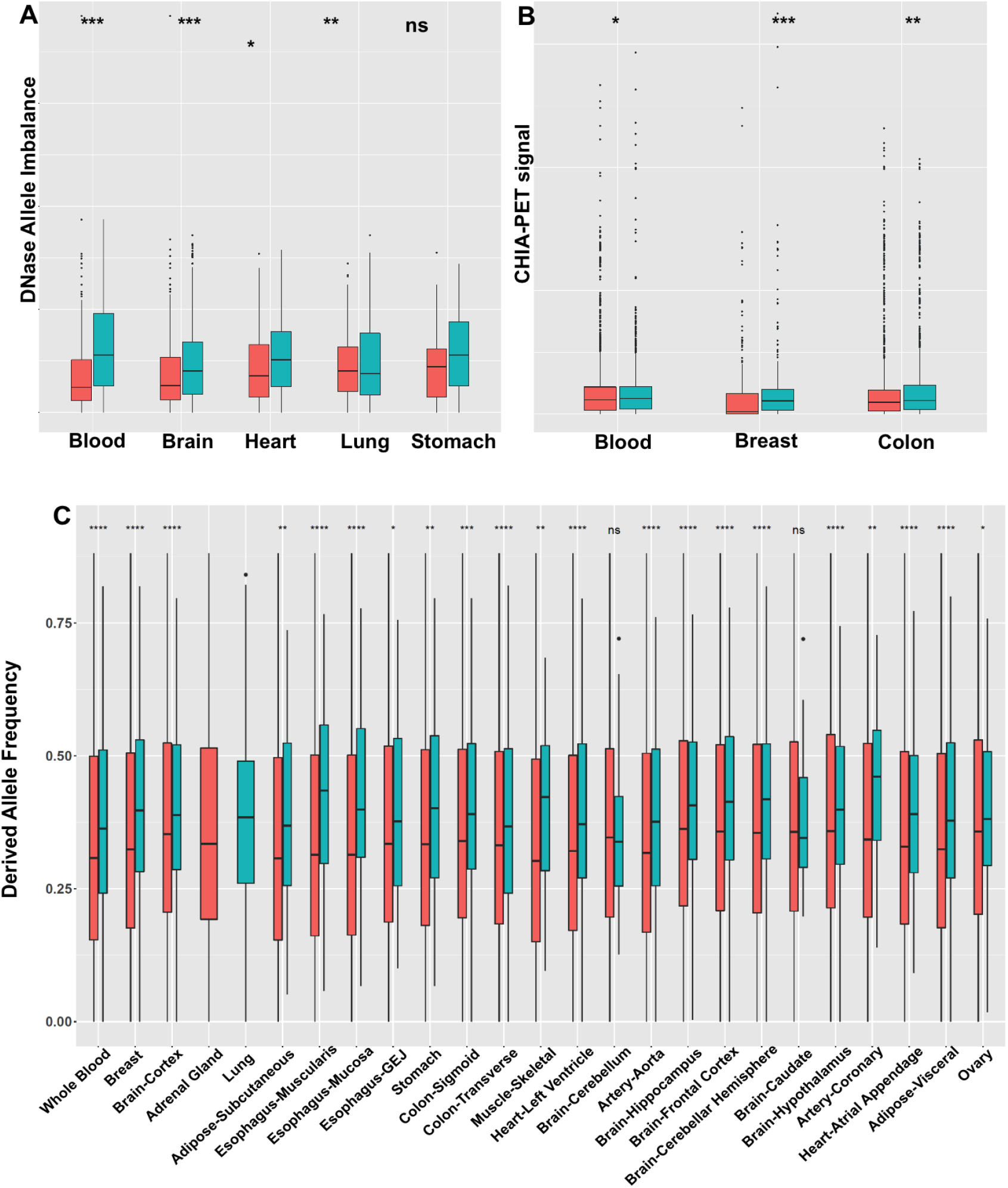
Detected SNPs exhibit allelic imbalance for open chromatin states, spatial proximity to target genes and higher derived allele frequency. (A) Detected SNPs exhibit significant allelic imbalance for open chromatin states. The foreground is the imbalance distribution of detected SNPs (blue), the control is the imbalance distribution generated from a null binomial model (red; see Methods). (B) Detected SNP-Gene pairs exhibit spatial proximity quantified by ChIA-PET data (pooled cell lines *K562, Hela, Nb4, and MCF7*). The foreground is detected SNP-Gene pairs (blue) and the control is random SNP-gene pairs within 1M bps, controlled to match the distance distribution of the foreground (red). The y-axis is the enrichment scores. (C) Detected SNPs (blue) exhibit significantly higher derived allele frequency compared to eSNPs (red) across 21 tissues out of 25.

To further assess functionality of our detected SNPs, we estimated allelic imbalance in DNA accessibility in five tissues (blood, brain, heart, lung, stomach). More specifically, for each tissue we obtained DNase-seq reads (merged from 2-3 samples as indicated in Supplementary Table 3) from ENCODE/RoadMap (Roadmap Epigenomics Consortium et al., 2015) and then performed variant calling (for detected age-interacting SNPs) to extract heterozygous sites for allele imbalance estimation (see Methods). The background was generated based on a binomial model with matched reads depth and equal probability of 0.5 for either allele. As shown in Fig. 5A, in 4 out of 5 tissues significantly higher-than-expected allelic imbalance were observed in our detected SNPs. Lack of clear signal in lung may be due to insufficient read depths in those specific DNase-seq samples.

Finally, we tested the tendency of detected SNP locus and the target gene locus to spatially co-localize in the nucleus, which would further attest to transcriptional regulation of the gene by the SNP locus. Since tissue-specific ChIA-PET data are relatively rare and of low resolution, we therefore followed (Das et al., 2015) and used ChIA-PET data merged from 4 cell lines *K562, Hela, Nb4 and MCF7*) from ENCODE, after which we quantified (Methods) and compared the spatial interaction of the detected SNP-Gene pairs with a background composed of interacting SNPs paired with randomly selected genes with the same distance distribution as the foreground. As shown in Fig. 5B, in three tissues (blood, breast, colon), the detected SNP regions exhibit significantly higher spatial proximity to their detected target genes than to the background (Wilcoxon test p-value = 4.7e-2, 1.8e-15, 9.3e-5). Overall, our results support, via several lines of evidence, potential functionality of the detected age-interacting SNPs and SNP-Gene pairs.

### Functional analyses of detected target genes and SNPs suggest links to age-related processes and complex diseases

The genes involved in our detected TF-SNP-Gene triplets are expected to exhibit age-associated expression in an allele-specific manner, which may have implication on ageing and age-associated complex diseases. To explore this further, first we performed functional enrichment analysis over target genes in a tissue-specific manner (see Methods). Fig. 6 shows a comprehensive TreeMap view of enriched GO terms across tissues (FDR <= 0.1; Methods) including GO terms that are significant enriched in at least 3 tissues. Enriched biological functions fall into twenty categories (Fig. 6). Among them, metabolic process (including molecular metabolism), cell death, DNA methylation, organelle organization (including cellular component assembly and telomere organization etc.), and negative regulation of biological process (including stress response, regulation of cell-cell communications etc.), have well-established links to aging (Licastro et al., 2005) and several age-associated complex diseases including hypertension, diabetes, neurodegeneration, and even cancer (Blasco et al., 2013; Jin, 2010; Sinclair et al., 2012). Metabolic process is crucial for the maintenance of homeostasis, which systematically deteriorates with aging (Barzilai et al., 2012). Programmed cell death is an important mechanism to maintain cellular hemostasis and eliminate pathological cells (Warner et al., 1997), which are both critical for aging process. DNA methylation is the most potent ageing biomarker and potentially could be related to tumorigenesis (Horvath, 2013). In addition, cellular stress response is a crucial biological process which modulates the damage to cells by activating repair signaling pathways. With aging, both cellular stress response and repair pathways decline, which could be a trigger for age-associated pathology.

**Fig. 6.**
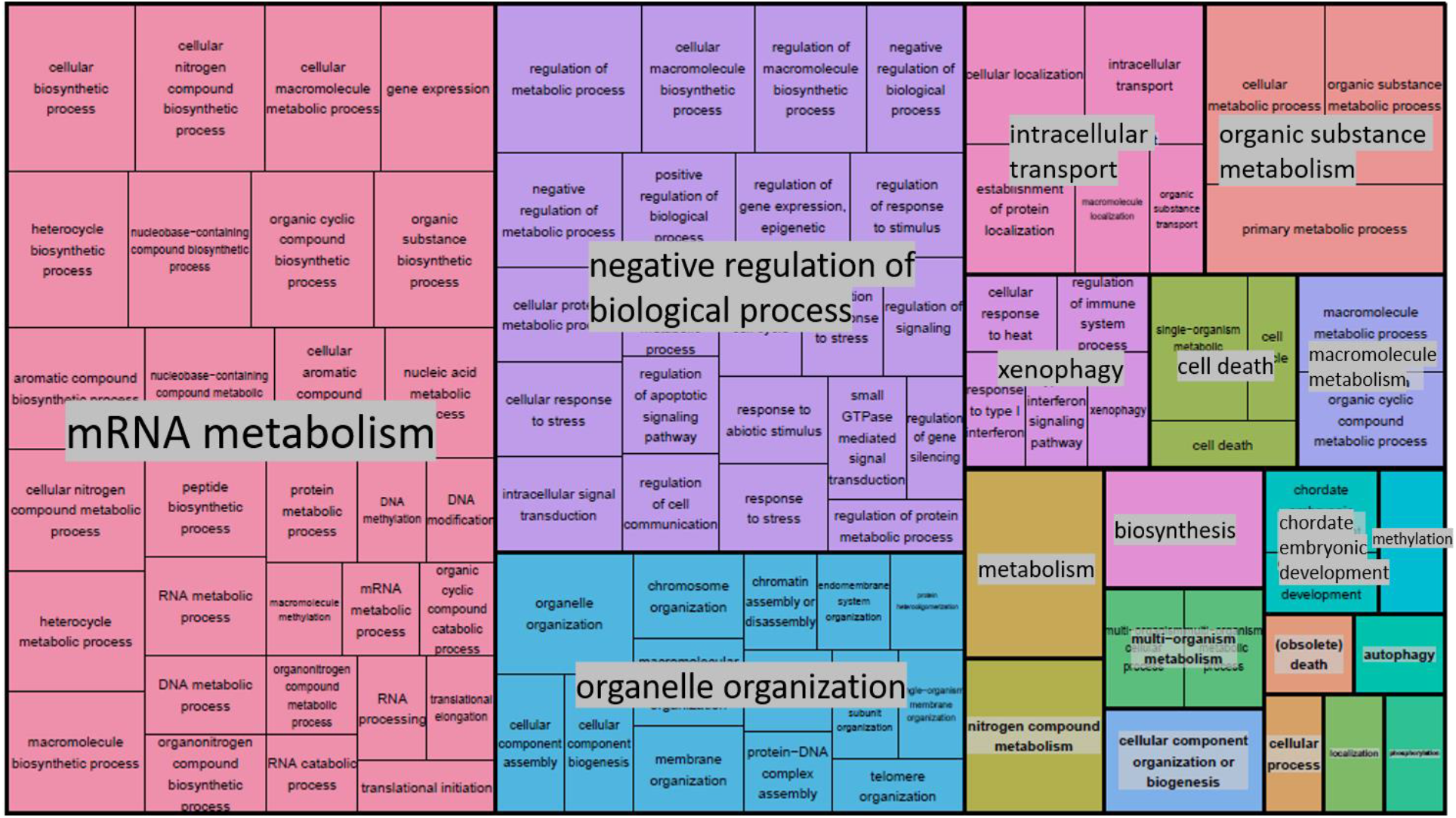
Enriched functions among the detected target genes. Enrichment analysis was done independently in each tissue, and the tree-map view shows the GO terms that were enriched (FDR < 5%) in at least three tissues.

Next, using Hypertension as an exemplar age-associated complex disease, we assessed the association of detected TF-SNP interactions with Hypertension in eight tissues (Adipose-Subcutaneous, Whole Blood, Artery - Aorta, Lung, Adipose-Visceral, Heart-Left Ventricle, Heart-Atrial Appendage, Artery-Coronary) that are implicated in hypertension. In each tissue independently, using log-likelihood-ratio (LLR) tests, we assessed whether aggregated TF-SNP interactions contribute to hypertension status in an individual (see Methods). As shown in Fig. 7, in 6 of the 8 tissues, aggregated interactions significantly contribute to hypertension.

**Fig. 7.**
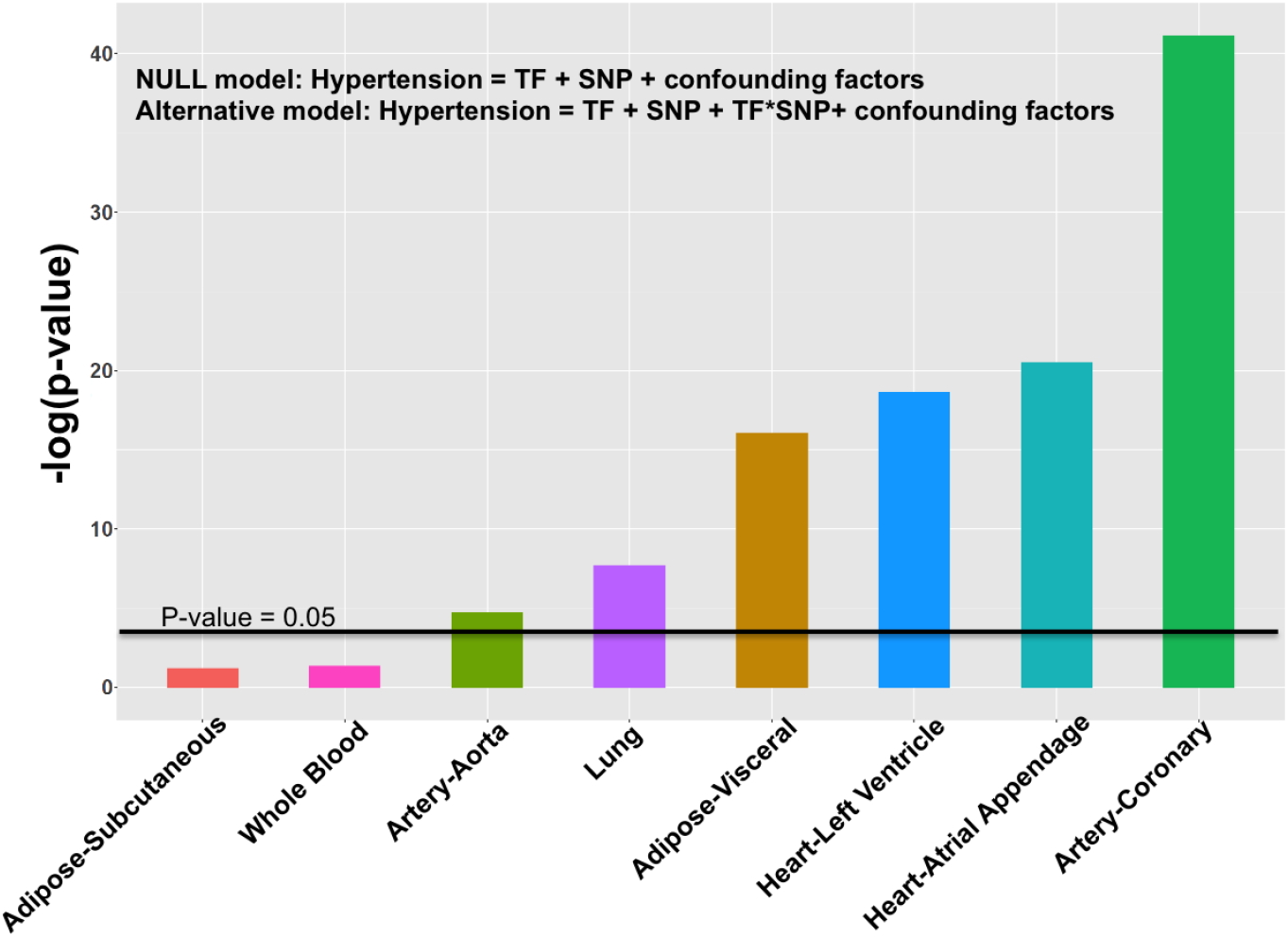
Detected TF-SNP interactions are potentially associated with Hypertension. Log likelihood ratio test was performed to assess the contribution of aggregated interactions to hypertension state of an individual in 8 tissues known to be etiologically linked to hypertension. The y axis is -Log(p-value of the log likelihood ratio test).

Previous studies have shown that tissue-specific eQTL SNPs (eSNP) exhibit significant overlap with those associated with various phenotypes. For instance, Higgins et al reported that eSNP rs43555985 (linked to gene GFRA2), which is shared with many other species, is associated with residual feed intake in beef cattle (Higgins et al., 2018). Luo et al reported that gene ZNF323 transcriptionally associated with GWAS SNPs rs1150711 and res2859365 is a potential schizophrenia causal candidate (Luo et al., 2015). We assessed whether the SNiPage-detected SNPs exhibit a similar, or greater, overlap with phenotype-associated SNPs. However, as an alternative to overlap statistics, we quantified and compared the distance of SNiPage-detected SNPs (as well as previously identified eSNPs as control) to phenotype-associated SNPs with the rationale that proximal SNPs are likely to be linked. We obtained GWAS signals for 8 diseases along with established corresponding tissue (Buniello et al., 2019; Lage et al., 2008). For each SNP (our detected SNPs as well as eSNPs), we measure its distance to the closest phenotype-associated SNP and compared the distances for the foreground and the eSNPs, using Wilcoxon test (see Methods). As shown in Fig. 8A, our detected SNPs are significantly closer to GWAS signals for breast cancer, cardiovascular diseases, colon cancer, and immune response disease. Even though it is not statistically significant for all the 8 diseases, the trend is still consistent in all cases.

**Fig. 8.**
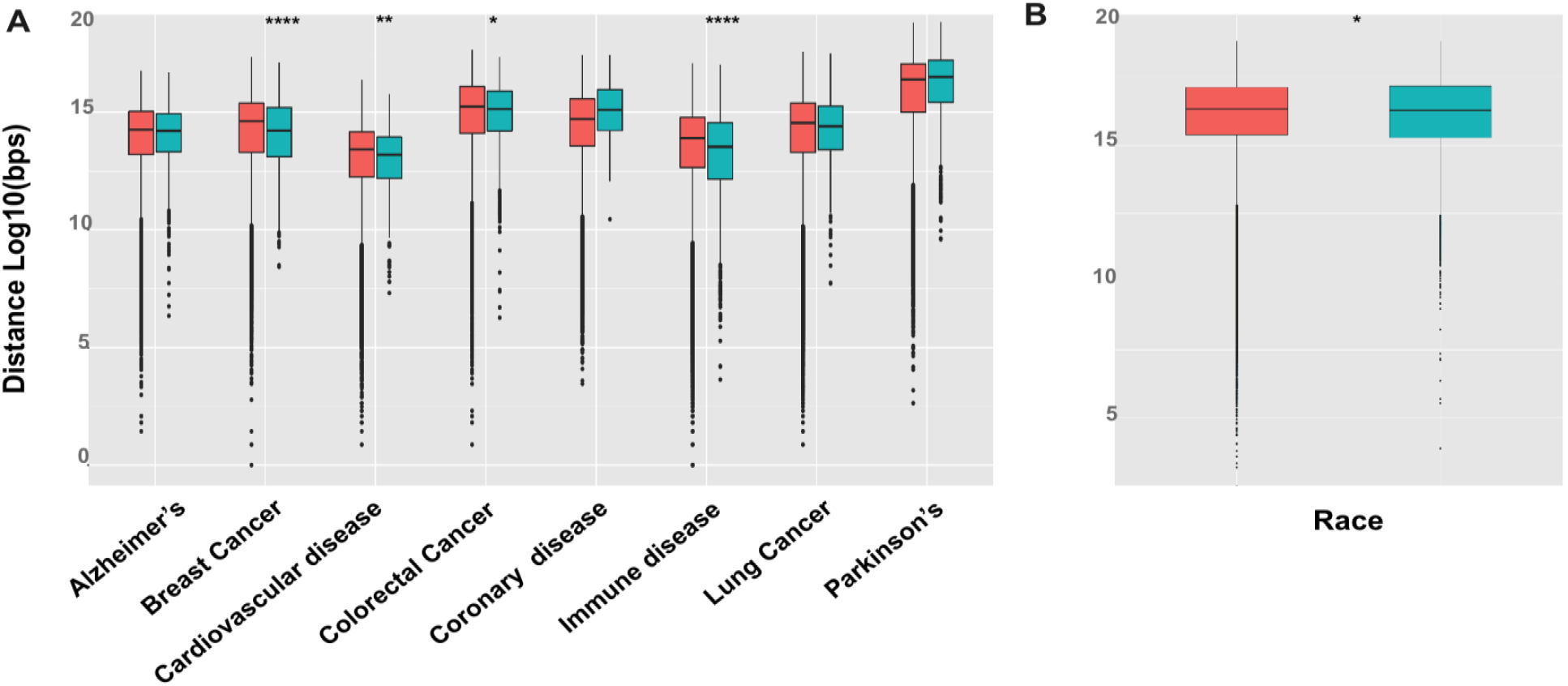
Detected SNPs are more likely to be associated with diseases and may reflect ethnicity structure. (A) Distribution of genomic distances of our detected SNPs (blue) and eSNPs (red) as control to the closest phenotype-associated SNPs. X-axis shows the phenotype. (B) Same as ‘A’ for known ethnicity-linked SNPs.

Additionally, we assessed whether our detected SNPs segregate with ethnicity. Interestingly, our detected SNPs, pooled across all tissues, exhibit greater genomic proximity to 299 ethnicity-associated SNPs (Huang et al., 2015) (Fig. 8B), suggesting that SNP-age interactions may partly explain the observed ethnicity-specific differences in age-associated diseases.

Taken together, our results suggest that target genes regulated by SNP-age interactions are potentially linked to crucial aging-related biological processes, aggregated interactions could potentially contribute to age-associated diseases such as hypertension, and relative to eSNPs, which are detected in an interaction-independent manner, SNiPage-detected SNPs are more likely to be causally linked to complex age-associated phenotypes.

## Discussion

We have reported a novel framework and a pipeline SNiPage, which incorporates a specific interaction mechanism to test the association between SNP-Age interactions and phenotypes (gene expression/diseases). We hypothesized that age-related TFs might preferentially bind to one of the alleles at a SNP, forming the basis for the interplay between the SNP and Age. We robustly detected numerous TF-SNP-Gene interaction triplets (average ~637) across 25 tissues. The detected SNP loci were enriched for epigenetic signals related to regulatory elements, exhibit allelic imbalance for chromatin accessibility, and exhibited spatial proximity to the putative target gene promoters, strongly supporting their functionality. Although our detected SNPs are under evolutionary conservation in human to the same extent as eSNPs, DAF analysis points to their evolution under positive (or relaxed purifying) selection during more recent human evolution, while at the same time, based on GO term enrichment and previously reported phenotype-associated SNPs, they are linked to aging and age-associated disease. This paradoxical result is, in fact, consistent with the “antagonistic pleiotropy” model of aging (Fabian, 2011), which posits that the dysfunction during aging may be a byproduct of allelic variants that are functionally adaptive during developmental and reproductive stages.

As noted above, although multiple previous studies incorporating intuitive SNP-Age terms in a regression framework have reported potential interactions (Simino et al., 2014; Yao et al., 2014), they suffer from both the detection power and insights about explicit interaction mechanisms, which are addressed by our approach. The numbers of SNiPage-detected TF-SNP interactions are orders of magnitude greater than the previously reported numbers of age-SNP interactions (~20). Using the age-associated TFs’ expression as proxy for age, and genome-wide gene expression values as molecular phenotype, allowed for detection of unprecedented numbers of significant SNP-Age interactions in multiple tissues. Furthermore, in contrast to previous studies, which identified interactions in a tissue-independent manner, our study identifies TF-SNP interactions independently in 25 tissues, enabling analysis of tissue-specific TF-SNP interaction effects on various phenotypes.

As an “environmental” risk factor, aging is likely to affect various age-associated phenotypes *via* gene regulation, which could be complex and diverse. Age-associated TFs represent just one specific agent of aging, and additional factors could be involved. For instance, age-associated splicing factor or epigenetic modification enzymes, both of which play essential roles in gene regulation and are linked to complex diseases, are potential agents of aging. We have previously observed age-associated splicing factor expression changes across tissues (Wang et al., 2018). Analogous to our current study, age-associated splicing factors can bind to polymorphic RNA motifs and form the basis for interaction between SNPs and aging. As for DNA methylation, Horvath et al. have shown methylation to be a robust biomarker of Age (Horvath, 2013) and moreover, it is well known that methylation usually blocks TF binding (Maurano et al., 2015; Moore et al., 2012; Yin et al., 2017). Thus, it is reasonable to expect that age-associated methylation changes could affect TF binding landscape across aging, in which case, age factor would interact with allele-specific binding sites to regulate gene expression. It is especially exciting to note that DNA methylation has also been identified as a robust biomarker for various cancers (Lakshminarasimhan and Liang, 2016). Combination of genotype, age-associated molecular changes including regulatory proteins and methylation, taken together, may provide novel insights into cancer.

Tissue-specificity of gene regulation complicates our understanding of complex diseases. We expect that our analysis across 25 tissues would shed light on molecular basis underlying tissue-specific genetic regulation. In fact, we specifically observed a greater number of interactions in tissues such as whole blood, Brain – Cortex and Colon - Transverse, which might imply that the genotype-associated functional diversity of those tissues may be more age-dependent. What’s more, our data can be used to draw links between tissues and complex diseases, as has been suggested based on eQTL studies (Acharya et al., 2017; Gilad et al., 2008). For instance, we found that in multiple known hypertension-related tissues our detected TF-SNP interactions significantly contribute, from statistical standpoint, to hypertension state of an individual.

Our results can inform the annotation and interpretation of GWAS and eQTL signals. In recent years, large scale GWAS and eQTL signals have been reported, however, it is still difficult to decode the observed associations into explicit pathways and mechanisms. Alignment of our detected interaction with reported associations by GWAS could suggest potential explicit links between environment, genetics, and phenotypes.

## Materials and Methods

### Transcriptome and Epigenomic data

We obtained the processed transcriptome data across 25 tissues from Genotype-Tissue Expression (GTEx) database version 6 (GTEx Consortium, 2015). Corresponding tissue-specific DNase-seq, broad peaks data for H3K27ac, H3K4me1 and H3K4me3 were downloaded from Roadmap consortium (Roadmap Epigenomics Consortium et al., 2015). In addition, top three principal components generated over SNP profile and PEER factors generated over gene expression profile were also obtained from GTEx database. GENCODE genome annotation version 19 (hg19) (Harrow et al., 2012) was used in this study.

### Detecting age-associated TFs

We used the linear regression model from previous studies (Glass et al., 2013; Yang et al., 2015) to detect significant age-associated genes as follows:

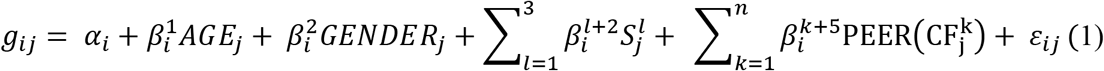

Where *g_ij_* is the expression of target gene i in jth sample and *α_i_* is the basal expression and the intercept on y axis for gene i. *β_i_* is the coefficient for covariates in the ith gene model. *AGE_j_* and *GENDER_j_* are the age and gender for jth sample respectively. *S_j_* is the covariate derived from SNPs for jth sample. 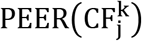 is the kth PEER factor (hidden variable) derived over gene expression profile for jth sample. *ε_ij_* denotes the error term.

We removed PEER factors, which significantly correlate with age (Pearson correlation coefficient < 0.05) (Brinkmeyer-Langford et al., 2016; Wang et al., 2018). TF genes for which the coefficient of age 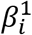 significantly deviated from zero (FDR <= 0.1) were considered age-associated.

### Putative allele-specific binding site prediction

We generated allele-specific sequence (~100 bps) around all the SNP loci and scanned the sequences for age-associated TFs’ binding sites based on published binding motifs in form of Positional Weight Matrix (PWM), using PWM-scan tool (Levy and Hannenhalli, 2002). Allele-specific binding is said to occur at a SNP when the putative TF binding predictions are significant (at least one PWM-SCAN hit with the p-value <= 0.005, which covers that SNP loci) in exactly one of the two alleles. SNPs having allele-specific binding were additionally filtered using tissue-specific DNase-seq broad peaks data to retain only the accessible loci to test for TF-SNP interactions.

### Significant Interaction detection

We modeled the association between the interaction (TF-SNP) and potential target gene expression (within 1 Mb) as follows:

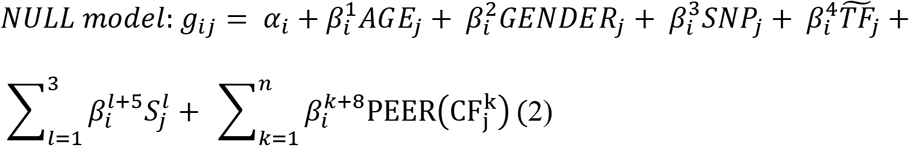

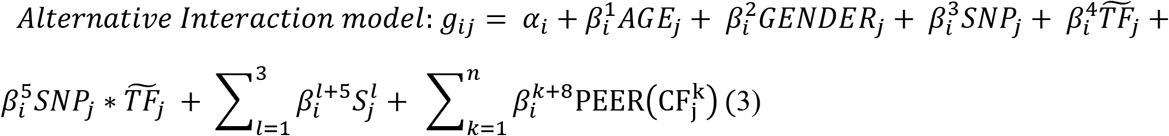

Equation (2) is the null model, and equation (3) is the alternative interaction model which includes the additional interaction term between SNP and TF. Log-likelihood-ratio tests were performed to evaluate the significance of interaction’s contribution to target gene expression. *g_ij_* is the expression of gene i in ith sample, *α_i_* is the basal expression and the intercept on y axis for gene i. *β_i_* is the coefficient for covariates in the ith gene model. *AGE_j_* and *GENDER_j_* are the age and gender respectively for jth sample. *SNP_j_* is allele frequency for ith sample. 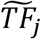 is adjusted concentration of TF which only includes age component (residuals after controlled for gender, covariates derived from SNP profile and PEER factors that are not correlated with age) for jth sample. The interaction term is denoted as 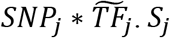 is the covariate derived from SNPs for jth sample and 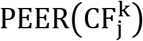 is the kth PEER factor (hidden variable) derived over gene expression profile for jth sample.

To select interactions consistent with our hypothesis and allele-specific TF binding prediction, we performed further filtering: we divided individuals into three categories: heterozygous (one binding and one non-binding allele), homozygous for binding allele, homozygous for non-binding allele. We retained the interactions only if the correlations between TF expression and target gene expression are significant for heterozygous and ‘homozygous for binding allele’ groups, but not for ‘homozygous for non-binding allele’ group. In addition, we require a greater correlation in ‘homozygous for binding allele group’ than for heterozygous group.

### Gene Functional analysis

We performed functional analysis on target genes involved in significant interaction across 25 tissues using R package “GoStats” (Falcon and Gentleman, 2007). Then a TreeMap view is generated using Revigo package (Supek et al., 2011) for the GO terms that were enriched (FDR <= 0.1) in at least 3 tissues.

### Enrichment test for Epigenomic signals

We extracted 100 bps windows centered at SNP loci and mapped raw DNase-seq, ChIP-seq signals for H3K27ac, H3K4me1, and H3K4me3 to those windows using “bedtools”. The average score across all the base pairs within the window is taken as the signal score. One tailed Wilcoxon test is performed to compare the signal score distributions between significant SNPs and other tested SNPs.

### Robustness test

We randomly under-sampled datasets (70%, 80%, 90%) to estimate the replication rate for detected interactions from the original dataset. The replicate rates were compared with that from the background dataset.

### DNase allelic imbalance test

We obtained 2-3 DNase-Seq samples (bam file after alignment) for each tissue and performed variant calling for our detected SNPs using “samtools” (Li et al., 2009). We specifically select heterozygous SNPs with at least 10 mapped reads. Then allelic imbalance score is estimated as |log2(reads from allele 1 + 1e-6 / reads from allele 2 + 1e-6)|. The background was sampled from a binomial distribution, which has the same read depth as the foreground and probabilities for observing either allele is 0.5. We estimated the allele imbalance for both foreground and background, then performed one tail Wilcoxon test to compare the two distributions.

### Hypertension association analysis

We model the association between interactions and hypertension as follows: The NULL model: Hypertension status ~ TF + SNP + confounding factors The interaction model: Hypertension status ~ TF + SNP + SNP*TF + confounding factors; The confounding factors used here are the same as ones in the interaction detection step. Log likelihood ratio test was performed to test whether detected SNP*TFs significantly contribute to hypertension. To reduce feature dimensionality, we performed PCA and took top 30 components to represent the information.

### CHIA-PET analysis

We obtained CHIA-Pet data from (Das et al., 2015), which merged the CHIA-Pet data from 4 cell lines (*K562, Hela, Nb4 and MCF7)*. Then we quantified the support for spatial proximity for a SNP-Gene pair using “bedtools”. To test the significance, we generate a background by selecting a random pair of SNP and target gene within 1M bp distance while controlling for the distance to match those for the foreground SNP-Gene pairs; for each pair of detected SNP and target gene, we selected a random pair of SNP and gene with approximate distance (within 10% difference) as corresponding background). We then applied one-sided wilcoxon test to compare the signal difference bwteen the foreground and the background.

### Conservation analysis

We obtained PhastCons scores derived from multiple alignment across 100 species from UCSC browser (Siepel et al., 2005). “bedtools” was used to calculate the mean PhastCons score across 100 bps centered around detected SNPs or background. We used three different background sets: random genome-wide SNPs, random SNPs that had passed DNase filtration, and tissue-specific eSNPs obtained from GTEx (GTEx Consortium, 2015). Wilcoxon test was used to compare the difference between foreground and background.

### Derived Allele Frequency

The variant calling files (VCF file) were obtained from 1000 genome consortium (Auton et al., 2015), then we calculated the DAF (derived allele frequency) for both our detected SNPs and tissue specific eSNPs from GTEx. Wilcoxon test was used to compare the difference.

### Phenotype association analysis for detected SNPs

We obtained 299 race-specific SNPs (Huang et al., 2015) and GWAS-identified SNPs for 8 diseases (Alzheimer’s diseases, breast cancer, colorectal cancer, lung cancer, cardiovascular diseases, coronary disease, Immune response disease, Parkinson’s disease). For each detected SNP, we calculated its distance to the closest phenotype-associated SNP. As a control, we applied the same procedure to eSNPs (SNPs detected by eQTL models) previously reported by GTEx (GTEx Consortium, 2015). One-sided Wilcoxon test was performed to compare the two distance distributions.

## Acknowledgment

Authors would like to thank Vishaka Datta and Mike Gertz for providing feedback on the initial draft. S.H. was funded in part by NSF award 1564785 to University of Maryland.

## Reference

Acharya, C. R., Owzar, K., Allen, A. S. and Carlo, M. (2017). Mapping eQTL by leveraging multiple tissues and DNA methylation. BMC Bioinformatics 1–11.

Adeyemo, A., Gerry, N., Chen, G., Herbert, A., Doumatey, A., Huang, H., Zhou, J., Lashley, K., Chen, Y., Christman, M., et al. (2009). A Genome-Wide Association Study of Hypertension and Blood Pressure in African Americans. 5, 1–11.

Auton, A., Abecasis, G. R., Altshuler, D. M., Durbin, R. M., Bentley, D. R., Chakravarti, A., Clark, A. G., Donnelly, P., Eichler, E. E., Flicek, P., et al. (2015). A global reference for human genetic variation. Nature 526, 68–74.

Barzilai, N., Huffman, D. M., Muzumdar, R. H. and Bartke, A. (2012). The Critical Role of Metabolic Pathways in Aging. Diabetes 61, 1315–1322.

Blasco, M. A., Partridge, L., Serrano, M., Kroemer, G. and Lo, C. (2013). Review The Hallmarks of Aging.

Brinkmeyer-Langford, C. L., Guan, J., Ji, G. and Cai, J. J. (2016). Aging Shapes the Population-Mean and -Dispersion of Gene Expression in Human Brains. Front. Aging Neurosci. 8, 183.

Buniello, A., Macarthur, J. A. L., Cerezo, M., Harris, L. W., Hayhurst, J., Malangone, C., McMahon, A., Morales, J., Mountjoy, E., Sollis, E., et al. (2019). The NHGRI-EBI GWAS Catalog of published genome-wide association studies, targeted arrays and summary statistics 2019. Nucleic Acids Res. 47, D1005–D1012.

Creyghton, M. P., Cheng, A. W., Welstead, G. G., Kooistra, T., Carey, B. W., Steine, E. J., Hanna, J., Lodato, M. A., Frampton, G. M., Sharp, P. A., et al. (2010). Histone H3K27ac separates active from poised enhancers and predicts developmental state. Proc. Natl. Acad. Sci. U. S. A. 107, 21931–6.

Das, A., Morley, M., Moravec, C. S., Tang, W. H. W., Hakonarson, H., Consortium, M., Margulies, K. B., Cappola, T. P., Jensen, S. and Hannenhalli, S. (2015). Bayesian integration of genetics and epigenetics detects causal regulatory SNPs underlying expression variability. Nat. Commun. 6, 1–11.

Dongen, J. Van, Nivard, M. G., Willemsen, G., Hottenga, J., Helmer, Q., Dolan, C. V, Ehli, E. A., Davies, G. E., Iterson, M. Van, Breeze, C. E., et al. (2016). Genetic and environmental influences interact with age and sex in shaping the human methylome. Nat. Commun. 7, 1–13.

Dror, H., Donyo, M., Atias, N., Mekahel, K., Melamed, Z., Yannai, S., Lev-Maor, G., Shilo, A., Schwartz, S., Barshack, I., et al. (2016). A network-based analysis of colon cancer Splicing changes reveals a tumorigenesis-favoring regulatory pathway emanating from ELK1. Genome Res. 26, 541–553.

Epigenetic regulation of RNA processing: Nature ENCODE: Nature Publishing Group. Fabian, D. K. (2014). The Evolution of Aging The Evolution of Aging Aging is an Evolutionary Paradox.

Falcon, S. and Gentleman, R. (2007). Using GOstats to test gene lists for GO term association. Bioinformatics 23, 257–258.

Gibney, E. R. and Nolan, C. M. (2010). Epigenetics and gene expression. Heredity (Edinb). 105, 4–13.

Gilad, Y., Rifkin, S. A. and Pritchard, J. K. (2008). Revealing the architecture of gene regulation: the promise of eQTL studies. Trends Genet. 24, 408–415.

Glass, D., Viñuela, A., Davies, M. N., Ramasamy, A., Parts, L., Knowles, D., Brown, A. A., Hedman, A. K., Small, K. S., Buil, A., et al. (2013). Gene expression changes with age in skin, adipose tissue, blood and brain. Genome Biol. 14, R75.

GTEx Consortium, Gte. (2015). Human genomics. The Genotype-Tissue Expression (GTEx) pilot analysis: multitissue gene regulation in humans. Science 348, 648–60.

Harrow, J., Frankish, A., Gonzalez, J. M., Tapanari, E., Diekhans, M., Kokocinski, F., Aken, B. L., Barrell, D., Zadissa, A., Searle, S., et al. (2012). GENCODE: The reference human genome annotation for the ENCODE project. Genome Res. 22, 1760–1774.

Higgins, M. G., Fitzsimons, C., McClure, M. C., McKenna, C., Conroy, S., Kenny, D. A., McGee, M., Waters, S. M. and Morris, D. W. (2018). GWAS and eQTL analysis identifies a SNP associated with both residual feed intake and GFRA2 expression in beef cattle. Sci. Rep. 8, 1–12.

Horvath, S. (2013). DNA methylation age of human tissues and cell types. Genome Biol. 14, R115.

Howard, A. D., Wang, X., Prasad, M., Sahu, A. Das, Aniba, R., Miller, M., Hannenhalli, S. and Chang, Y. P. C. (2019). Allele-specific enhancers mediate associations between LCAT and ABCA1 polymorphisms and HDL metabolism. PLoS One 14, 1–17.

Huang, T., Shu, Y. and Cai, Y. D. (2015). Genetic differences among ethnic groups. BMC Genomics 16, 1–10.

Jiang, Y., Shen, H., Liu, X., Dai, J., Jin, G., Qin, Z., Chen, J., Wang, S., Wang, X., Hu, Z., et al. (2011). Genetic variants at 1p11.2 and breast cancer risk: A two-stage study in Chinese women. PLoS One 6,.

Jin, K. (2010). Modern Biological Theories of Aging. 1, 72–74.

Lage, K., Hansen, N. T., Karlberg, E. O., Eklund, A. C., Roque, F. S., Donahoe, P. K., Szallasi, Z., Jensen, T. S. and Brunak, S. (2008). A large-scale analysis of tissue-specific pathology and gene expression of human disease genes and complexes. Proc. Natl. Acad. Sci. 105, 20870–20875.

Lakshminarasimhan, R. and Liang, G. (2016). The role of DNA methylation in cancer. Adv. Exp. Med. Biol. 945, 151–172.

Levy, S. and Hannenhalli, S. (2002). Identification of transcription factor binding sites in the human genome sequence. 514, 510–514.

Li, H., Handsaker, B., Wysoker, A., Fennell, T., Ruan, J., Homer, N., Marth, G., Abecasis, G. and Durbin, R. (2009). The Sequence Alignment/Map format and SAMtools. Bioinformatics 25, 2078–2079.

Licastro, F., Candore, G., Lio, D., Porcellini, E., Colonna-romano, G., Franceschi, C. and Caruso, C. (2005). Innate immunity and inflammation in ageing: a key for understanding age-related diseases. Immun. Aging 14, 1–14.

Lubbe, S. J., Pittman, a M., Olver, B., Lloyd, a, Vijayakrishnan, J., Naranjo, S., Dobbins, S., Broderick, P., Gómez-Skarmeta, J. L. and Houlston, R. S. (2012). The 14q22.2 colorectal cancer variant rs4444235 shows cis-acting regulation of BMP4. Oncogene 31, 3777–84.

Luco, R. F., Pan, Q., Tominaga, K., Blencowe, B. J., Pereira-Smith, O. M. and Misteli, T. (2010). Regulation of alternative splicing by histone modifications. Science 327, 996–1000.

Luo, X. J., Mattheisen, M., Li, M., Huang, L., Rietschel, M., Børglum, A. D., Als, T. D., Van Den Oord, E. J., Aberg, K. A., Mors, O., et al. (2015). Systematic Integration of Brain eQTL and GWAS Identifies ZNF323 as a Novel Schizophrenia Risk Gene and Suggests Recent Positive Selection Based on Compensatory Advantage on Pulmonary Function. Schizophr. Bull. 41, 1294–1308.

Maurano, M. T., Wang, H., John, S., Canfield, T., Lee, K. and Stamatoyannopoulos, J. A. (2015). Role of DNA Methylation in Modulating Transcription Article Role of DNA Methylation in Modulating Transcription Factor Occupancy. CellReports 12, 1184–1195.

Monlong, J., Calvo, M., Ferreira, P. G. and Guigo, R. (2014). Identification of genetic variants associated with alternative splicing using sQTLseekeR. Nat. Commun.

Moore, L. D., Le, T. and Fan, G. (2012). DNA Methylation and Its Basic Function. Neuropsychopharmacology 38, 23–38.

Niccoli, T. and Partridge, L. (2012). Ageing as a Risk Factor for Disease. Curr. Biol. 22, R741–R752.

Rantalainen, M., Lindgren, C. M. and Holmes, C. C. (2015). Robust Linear Models for Cis-eQTL Analysis. PLoS One 1–16.

Roadmap Epigenomics Consortium, Kundaje, A., Meuleman, W., Ernst, J., Bilenky, M., Yen, A., Heravi-moussavi, A., Kheradpour, P., Zhang, Z., Wang, J., et al. (2015). Integrative analysis of 111 reference human epigenomes. Nature 317–330.

Siepel, A., Bejerano, G., Pedersen, J. S., Hinrichs, A. S., Hou, M., Rosenbloom, K., Clawson, H., Spieth, J., Hillier, L. D. W., Richards, S., et al. (2005). Evolutionarily conserved elements in vertebrate, insect, worm, and yeast genomes. Genome Res. 15, 1034–1050.

Simino, J., Shi, G., Bis, J. C., Chasman, D. I., Ehret, G. B., Lyytika, L., Nolte, I. M., Sim, X., Dehghan, A., Eiriksdottir, G., et al. (2014). Gene-Age Interactions in Blood Pressure Regulation: A Large-Scale Investigation with the CHARGE, Global BPgen, and ICBP Consortia. Am. J. Hum. Genet. 24–38.

Sinclair, D., North, B., Editors, G., North, B. J. and Sinclair, D. A. (2012). The Intersection Between Aging and Cardiovascular Disease. 02115, 1097–1108.

Sug, S., Yoon, S., Ph, D., Fryar, C. D. and Carroll, M. D. (2015). Hypertension Prevalence and Control Among Adults: United States, 2011-2014. 2011-2014.

Supek, F., Bošnjak, M., Škunca, N. and Šmuc, T. (2011). Revigo summarizes and visualizes long lists of gene ontology terms. PLoS One 6,.

Tollervey, J. R., Wang, Z., Hortobágyi, T., Witten, J. T., Zarnack, K., Kayikci, M., Clark, T. A., Schweitzer, A. C., Rot, G., Curk, T., et al. (2011). Analysis of alternative splicing associated with aging and neurodegeneration in the human brain. Genome Res. 21, 15721582.

Visscher, P. M., Wray, N. R., Zhang, Q., Sklar, P., Mccarthy, M. I., Brown, M. A. and Yang, J. (2017). 10 Years of GWAS Discovery: Biology, Function, and Translation. Am. J. Hum. Genet. 101, 5–22.

Wang, K., Das, A., Xiong, Z., Cao, K. and Hannenhalli, S. (2014a). Phenotype-dependent coexpression gene clusters: application to normal and premature ageing. IEEE/ACM Trans. Comput. Biol. Bioinforma. 1–1.

Wang, H., Burnett, T., Kono, S., Haiman, C. A., Iwasaki, M., Wilkens, L. R., Loo, L. W. M., Berg, D. Van Den, Kolonel, L. N., Henderson, B. E., et al. (2014b). Trans-ethnic genomewide association study of colorectal cancer identifies a new susceptibility locus in VTI1A. Nat. Commun. 1–7.

Wang, K., Wu, D., Zhang, H., Das, A., Basu, M., Malin, J., Cao, K. and Hannenhalli, S. (2018). Comprehensive map of age-associated splicing changes across human tissues and their contributions to age-associated diseases. Sci. Rep. 8, 10929.

Warner, H. R., Hodes, R. J. and Pocinki, K. (1997). What does cell death have to do with aging? J. Am. Geriatr. Soc. 45, 1140–1146.

Westra, H. and Franke, L. (2014). From genome to function by studying eQTLs. BBA - Mol. Basis Dis. 1842, 1896–1902.

Wyss-coray, T. (2015). Ageing, neurodegeneration and brain rejuvenation.

Yang, J., Huang, T., Petralia, F., Long, Q., Zhang, B., Argmann, C., Zhao, Y., Mobbs, C. V, Schadt, E. E., Zhu, J., et al. (2015). Synchronized age-related gene expression changes across multiple tissues in human and the link to complex diseases. Sci. Rep. 5, 15145.

Yao, C., Joehanes, R., Johnson, A. D., Huan, T., Ying, S., Freedman, J. E., Murabito, J., Lunetta, K. L., Metspalu, A., Munson, P. J., et al. (2014). Sex- and age-interacting eQTLs in human complex diseases. Hum. Mol. Genet. 23, 1947–1956.

Yin, Y., Morgunova, E., Jolma, A., Kaasinen, E., Sahu, B., Khund-sayeed, S., Das, P. K., Kivioja, T., Dave, K., Zhong, F., et al. (2017). Impact of cytosine methylation on DNA binding specificities of human transcription factors. 2239,.

Zhao, K., Lu, Z., Park, J. W., Zhou, Q. and Xing, Y. (2013). GLiMMPS: robust statistical model for regulatory variation of alternative splicing using RNA-seq data. Genome Biol. 14, R74.

