## Supplemental Figures and Tables for "A transcription-centric model of SNP-Age interaction"

### Supplementary Figures

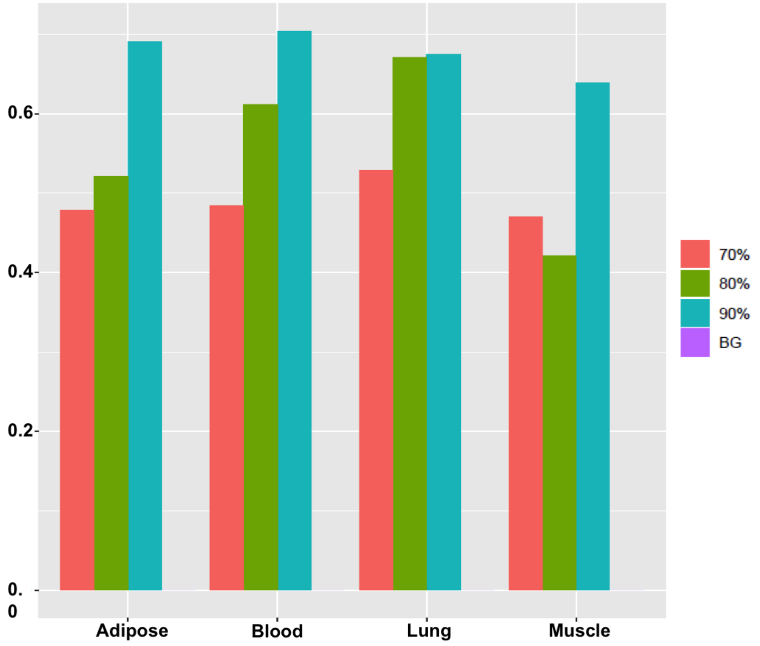


**Supplementary Figure 1: Replication rates for under-sampling (70%, 80%, 90%) across 4 tissues (Robustness analysis).**

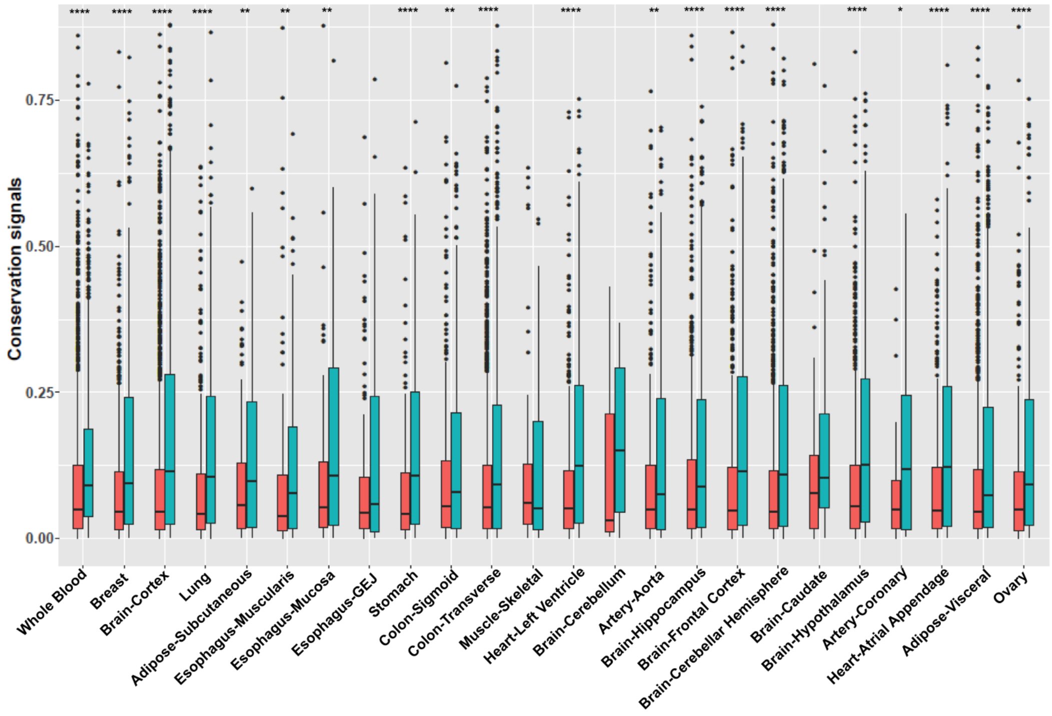


**Supplementary Figure 2: Average conservation score for detected SNPs and random SNPs (without passing DNase filtration) across tissues.**

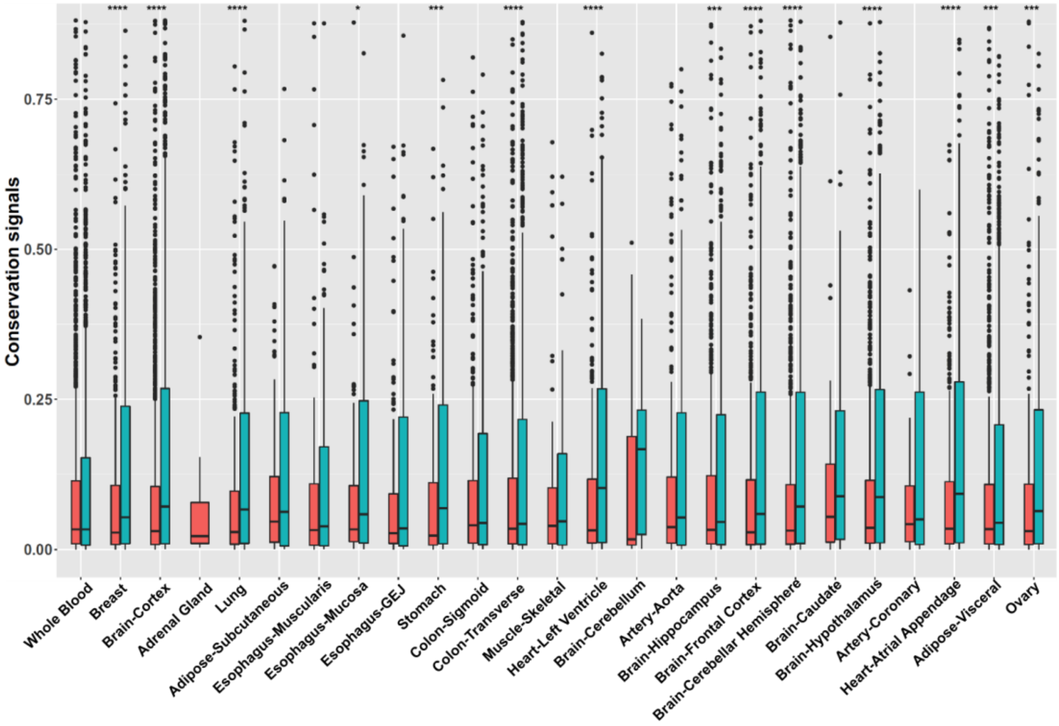


**Supplementary Figure 3: Average conservation score for detected SNPs and random SNPs (after passing DNase filtration) across tissues.**

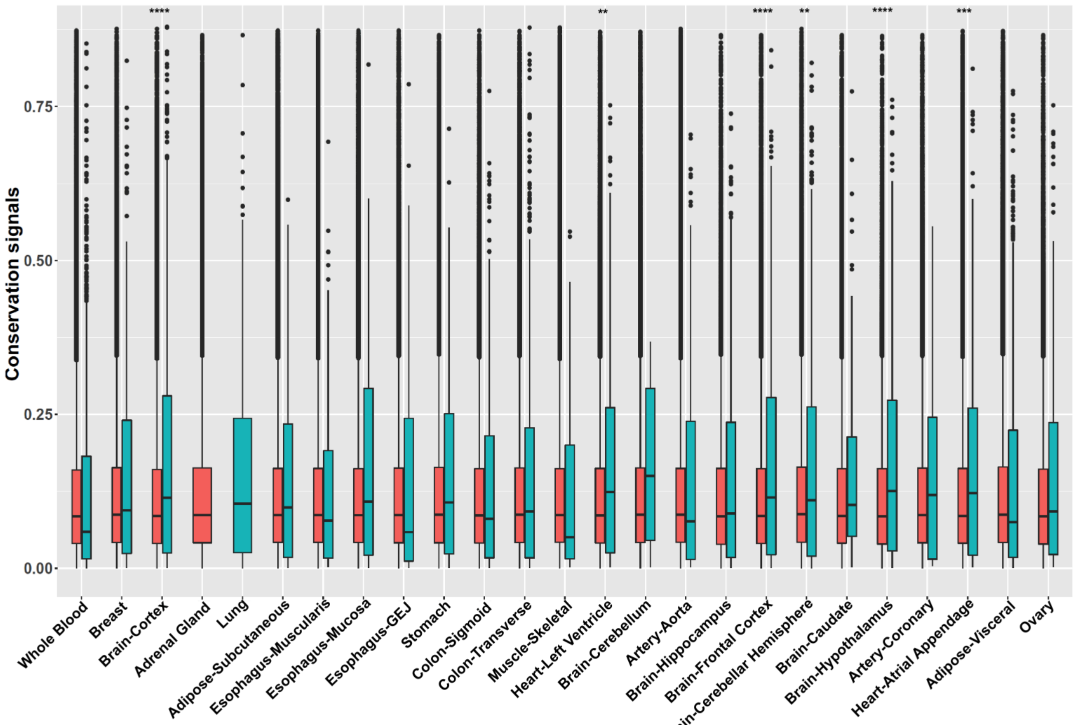


**Supplementary Figure 4: Average conservation score for detected SNPs and eSNPs across tissues.**

### Supplementary Tables

#### Supplementary table 1:

| Tissues | Number of samples |
|---|---|
| Whole Blood | 338 |
| Adipose-Subcutaneous | 298 |
| Muscle-Skeletal | 361 |
| Artery-Coronary | 118 |
| Heart-Atrial Appendage | 159 |
| Adipose-Visceral | 185 |
| Ovary | 85 |
| Breast-Mammary | 183 |
| Brain-Cortex | 96 |
| Adrenal Gland | 126 |
| Lung | 278 |
| Esophagus-Muscularis | 218 |
| Esophagus-Mucosa | 241 |
| Esophagus-Gastroesophageal Junction | 127 |
| Stomach | 170 |
| Colon-Sigmoid | 124 |
| Colon-Transverse | 169 |
| Heart-Left Ventricle | 190 |
| Brain-Cerebellum | 103 |
| Artery-Aorta | 197 |
| Brain-Hippocampus | 81 |
| Brain-Frontal Cortex | 92 |
| Brain-Cerebellar Hemisphere | 89 |
| Brain-Caudate | 100 |
| Brain-Hypothalamua | 81 |

**Supplementary table 2:**

| Tissues | Number of triplets | Number of target genes | Number of TFs | Number of SNPs |
|---|---|---|---|---|
| Whole Blood | 70 | 65 | 52 | 65 |
| Adipose-Subcutaneous | 28 | 23 | 20 | 28 |
| Muscle-Skeletal | 11 | 10 | 9 | 11 |
| Artery-Coronary | 10 | 9 | 8 | 10 |
| Heart-Atrial Appendage | 129 | 120 | 68 | 122 |
| Adipose-Visceral | 238 | 221 | 119 | 221 |
| Ovary | 194 | 190 | 82 | 188 |
| Breast-Mammary | 210 | 196 | 96 | 196 |
| Brain-Cortex | 259 | 243 | 106 | 243 |
| Adrenal Gland | 22 | 22 | 14 | 21 |
| Lung | 65 | 62 | 51 | 64 |
| Esophagus-Muscularis | 30 | 30 | 21 | 29 |
| Esophagus-Mucosa | 45 | 43 | 35 | 44 |
| Esophagus-Gastroesophageal Junction | 101 | 96 | 57 | 97 |
| Stomach | 205 | 192 | 54 | 188 |
| Colon-Sigmoid | 103 | 98 | 70 | 100 |
| Colon-Transverse | 227 | 201 | 86 | 189 |
| Heart-Left Ventricle | 31 | 28 | 26 | 28 |
| Brain-Cerebellum | 9 | 9 | 4 | 9 |
| Artery-Aorta | 142 | 137 | 80 | 133 |
| Brain-Hippocampus | 149 | 141 | 48 | 140 |
| Brain-Frontal Cortex | 125 | 119 | 33 | 114 |
| Brain-Cerebellar Hemisphere | 178 | 166 | 69 | 169 |
| Brain-Caudate | 14 | 14 | 7 | 14 |
| Brain-Hypothalamua | 162 | 154 | 62 | 153 |
